# The meditative brain: State and trait changes in harmonic complexity for long-term mindfulness meditators

**DOI:** 10.1101/2023.11.16.567347

**Authors:** Selen Atasoy, Anira Escrichs, Eloise Stark, Kendra G. M. Terry, Estela Camara, Ana Sanjuan, Shamil Chandaria, Gustavo Deco, Morten L. Kringelbach

**Affiliations:** Centre for Eudaimonia and Human Flourishing, University of Oxford, Oxford, United Kingdom; Center of Music in the Brain (MIB), Clinical Medicine, Aarhus University; Computational Neuroscience Group, Center for Brain and Cognition, Department of Information and Communication Technologies, Universitat Pompeu Fabra, Barcelona, Catalonia, Spain; Teachers College, Columbia University, New York, USA; Cognition and Brain Plasticity Unit, Bellvitge Biomedical Research Institute (IDIBELL), L’Hospitalet de Llobregat, Barcelona, Spain; Department of Cognition, Development and Educational Psychology, University of Barcelona, Barcelona, Spain; ICREA, Institució Catalana de Recerca i Estudis Avancats (ICREA), Spain; Department of Neuropsychology, Max Planck Institute for Human Cognitive and Brain Sciences, Leipzig, Germany; School of Psychological Sciences, Monash University, Melbourne, Australia

## Abstract

Meditation is an ancient practice that is shown to yield benefits for cognition, emotion regulation and human flourishing. In the last two decades, there has been a surge of interest in extracting the neural correlates of meditation, in particular of mindfulness meditation. Yet, these efforts have been mostly limited to the analysis of certain regions or networks of interest and a clear understanding of meditation-induced changes in the whole-brain dynamics has been lacking. Here, we investigate meditation-induced changes in brain dynamics using a novel connectome-specific harmonic decomposition method. Specifically, utilising the connectome harmonics as brain states - elementary building blocks of complex brain dynamics - we study the immediate (state) and long-term (trait) effects of mindfulness meditation in terms of the energy, power and complexity of the repertoire of these harmonic brain states. Our results reveal increased power, energy and complexity of the connectome harmonic repertoire and demonstrate that meditation alters brain dynamics in a frequency selective manner. Remarkably, the frequency-specific alterations observed in meditation are reversed in resting state in group-wise comparison revealing for the first time the long-term (trait) changes induced by meditation. These findings also provide evidence for the entropic brain hypothesis in meditation and provide a novel understanding of state and trait changes in brain dynamics induced by mindfulness meditation revealing the unique connectome harmonic signatures of the meditative brain.

## Introduction

Originating from Ancient Eastern traditions, meditation refers to a set of mental training practices developed for cultivating attentional and emotional self-regulation, well-being and emotional balance^1,2^. Based on the primary cognitive mechanisms involved, three distinct families of meditation have been identified: (1) attentional, (2) constructive, and (3) deconstructive families with primary cognitive mechanisms of (1) attention regulation and meta-awareness, (2) perspective taking and reappraisal and (3) self-inquiry, respectively^3^.

Attentional family meditation involves focusing and training various processes related to attention regulation. In this family, focused attention meditation (FOM) and open monitoring meditation (OMM) are the most commonly studied^2^. In FOM, the practitioner focuses their attention on a chosen object such as the breath. During OMM, the attention is directed to openly monitoring the experience from moment to moment, without reacting to its content. Although OMM is similar to deconstructive family meditations, it differs from this family by its primary objective, which is the stabilization of meta-awareness in relation to a particular attentional configuration^3^.

The constructive family involves the meditation practices supporting psychological patterns, which foster well-being by replacing maladaptive self-beliefs with positive self-conception such as loving-kindness and compassion^3^.

In the deconstructive family, a similar attentional focus on experience is maintained, however the purpose is to cultivate insight into one’s perspective of self, and the world. Therefore, this family focusses on self-inquiry, i.e., the process of investigating the nature of conscious experience^3^. Mindfulness (insight) meditation is the most well-known example of this family of meditation. Perhaps confusingly, mindfulness is sometimes associated with OMM as well. However, this is due to broad use of the word mindfulness since the development of its clinical applications such as mindfulness-based stress reduction (MBSR) and mindfulness-based cognitive therapy (MBCT). Classically, mindfulness refers to insight (Vipassanā) meditation, which involves inquiring into the nature of experience.

In neuroscientific literature, mindfulness meditation, has received significant attention over the last decades. Mindfulness has been defined as “the awareness that emerges through paying attention on purpose, in the present moment, and non-judgmentally to the unfolding of experience moment by moment”^4^. It has been described as “inherently a state of consciousness”^5^.

In mindfulness meditation, which originates from Vipassanā meditation, one focuses and re-focuses awareness on the present moment. The object of focus could be either internal or external experiences, such as breathing, thoughts, ambient sounds, or interoceptive sensations. The meditative brain state does not involve active problem solving, and aims for limited mind-wandering. In Vipassanā meditation, which follows the “open monitoring” style, training includes attention to the transient nature of sensory experience and shifting of attention across sensory modalities, with a non-judgemental attitude^6^. Mindfulness meditation also often requires re-perceiving, which is the capacity to witness the contents of one’s consciousness with no or little emotional and cognitive reactivity^7^. Deikman^8^ describes this state as a strengthening of the observing self.

Significant research over the past three decades has explored the efficacy of mindfulness meditation and clinical interventions using mindfulness such MBSR and MBCT. Over a hundred randomized controlled trials (RCTs) are listed on PubMed, with a firm majority demonstrating a benefit of mindfulness compared to active controls in both mental and physical health conditions, as well as in healthy individuals^9^. Meta-analyses have confirmed positive effects of mindfulness meditation on common mental health symptoms, including anxiety, depression and stress^9^. It should be noted that MBSR has been found to be more effective than ‘pure’ mindfulness meditation in some cases, possibly due to additional psychoeducational components or non-specific therapeutic contact^10^.

From a neuroscientific perspective, a fundamental question surrounding meditation is how it exerts the positive benefits of attentional and emotional self-regulation^11^ and resilience against recurrent depression^12^. Over the last decade, there has been a surge of interest in extracting the neural correlates of meditation^1,13–15^ and in particular mindfulness meditation mostly driven by its positive clinical implications. While early studies focused on extracting the neural correlates in terms of increased or decreased activity of certain brain regions using region-of-interest (ROI) analysis^12,16,17^, later studies involved more complex network based connectivity analysis^1,13–15,18–21^.

In ROI-based analysis, mindfulness meditation is found to have increased activity in the brain areas involved in autobiographical memory, self-reference processing and in the limbic system (right basal ganglia, right insula, right subgenual ACC, gyrus rectus, ventromedial prefrontal cortex, right ventrolateral prefrontal cortex (vlPFC), and right superior frontal gyrus)^16^, and in posterior cingulate cortex (PCC) and the dorsomedial prefrontal cortex (dmPFC)^17^, which are cortical regions associated with higher order cognitive and emotional functions, including meta-awareness, self-referencing and “thinking about thinking^22^. Network-based connectivity analysis has revealed changes in the connectivity of one particular network, the default-mode network (DMN), involved in mind-wandering and self-referential processing^1,13,18^. Alterations in DMN functional connectivity have been also found in focused attention meditation (FAM) and open monitoring meditation (OMM)^15^. Remarkably, mindfulness mediation has been also found to induce changes in structural connectivity in insula networks^19^.

Recently, in an attempt to provide insights into brain dynamics altered by meditation, some studies have focused on inter-network connectivity^14^ and dynamic functional connectivity^21^. Studying inter-network connectivity revealed increased functional connectivity between the DMN and the dorsal attention network (DAN), indicating meditation enables fast switching between mind wandering and focused attention states and better maintenance of attention once in the attentive state^14^. Studies using dynamic functional connectivity - a method to characterize functional connectivity changes over time - have found that participants with higher trait mindfulness scores showed more frequent switching between identified brain connectivity states^21^, which has been also replicated in children and adolescents^23^ and also spent significantly more time in a brain connectivity state associated with task-readiness^21^ and focused attention^20^. More recently, whole-brain computational modelling has also been utilized to understand the short- and long-term effects of mindfulness meditation^24–26^. Although these studies provide valuable insights into alterations of brain’s functional networks by meditation, they are all based on a priori definition of brain’s functional network architecture. Yet a recent study has shown that meditation not only changes the amount of functional connectivity within and between brain’s networks but also leads to alterations in the boundaries and configurations of these networks suggesting a reconfiguration whole-brain network architecture^27^.

In this work, we study the short and long-term effects of mindfulness meditation on brain dynamics. To this end, we extract the dynamical changes in cortical activity occurring during focused attention on breathing meditation and resting state in two different groups, i.e. experienced Vipassanā meditators and meditation-naive healthy controls using the novel framework of connectome harmonic decomposition.

## Results

### Connectome harmonic decomposition of fMRI data

Connectome harmonic decomposition (CHD) studies brain activity measured by the (fMRI BOLD signal) by decomposing it into a set of harmonic brain states defined by connectome harmonics^28,29^. By definition connectome harmonics extend the well-known Fourier transform to the particular structural connectivity of the human brain (i.e. the human connectome)^30,31^. The same way the Fourier transform decomposes any signal into a combination of sine and cosines, CHD enables the decomposition of any pattern of brain activity into the combination of connectome-specific harmonic waves. Thus, CHD represents brain activity in a new frequency-specific harmonic lanfuage and allows for the study of complex brain dynamics in terms of activation of elementary brain states, i.e. fundamental building blocks of brain activity^29^. This framework has been successfully applied to decode neural signatures of various states of consciousness including task-based brain activity in healthy population^32^, psychedelic-induced altered state of consciousness^28,33^, propofol-induced loss of consciousness^31^ and vegetative state and minimally conscious state patients^31^. Here, we apply the framework of CHD for the first time to extract the harmonic signatures underlying the short-term and long-term effects of mindfulness meditation.

Firstly, we estimate the connectome harmonic patterns as introduced in^30^ in a group averaged manner as in^28^ (Figure 1). To this end, T1 magnetic resonance imaging (MRI) data is used to reconstruct the cortical surface between gray and white matter. Diffusion tensor imaging (DTI) data is used to extract thalamo-cortical fibers using deterministic fiber tractography. Both, local and long-distance connections are combined to create the connectivity matrix of the human connectome (Figure 1**b**). Connectome harmonics 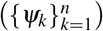 are estimated by applying the Laplace operator ∆ to the group averaged human connectome and computing its eigenvectors (∆*ψ*_*k*_ = *λ*_*k*_*ψ*_*k*_) (Figure 1**c**). Functional magnetic resonance imaging (fMRI) data (*ℱ*) as illustrated in (Figure 1 **d**) is decomposed in to the activation of connectome harmonics 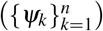 yielding the power of activation of each of these brain states for each time instance 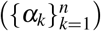 as delineated in (Figure 1 **e**).

**Figure 1.**
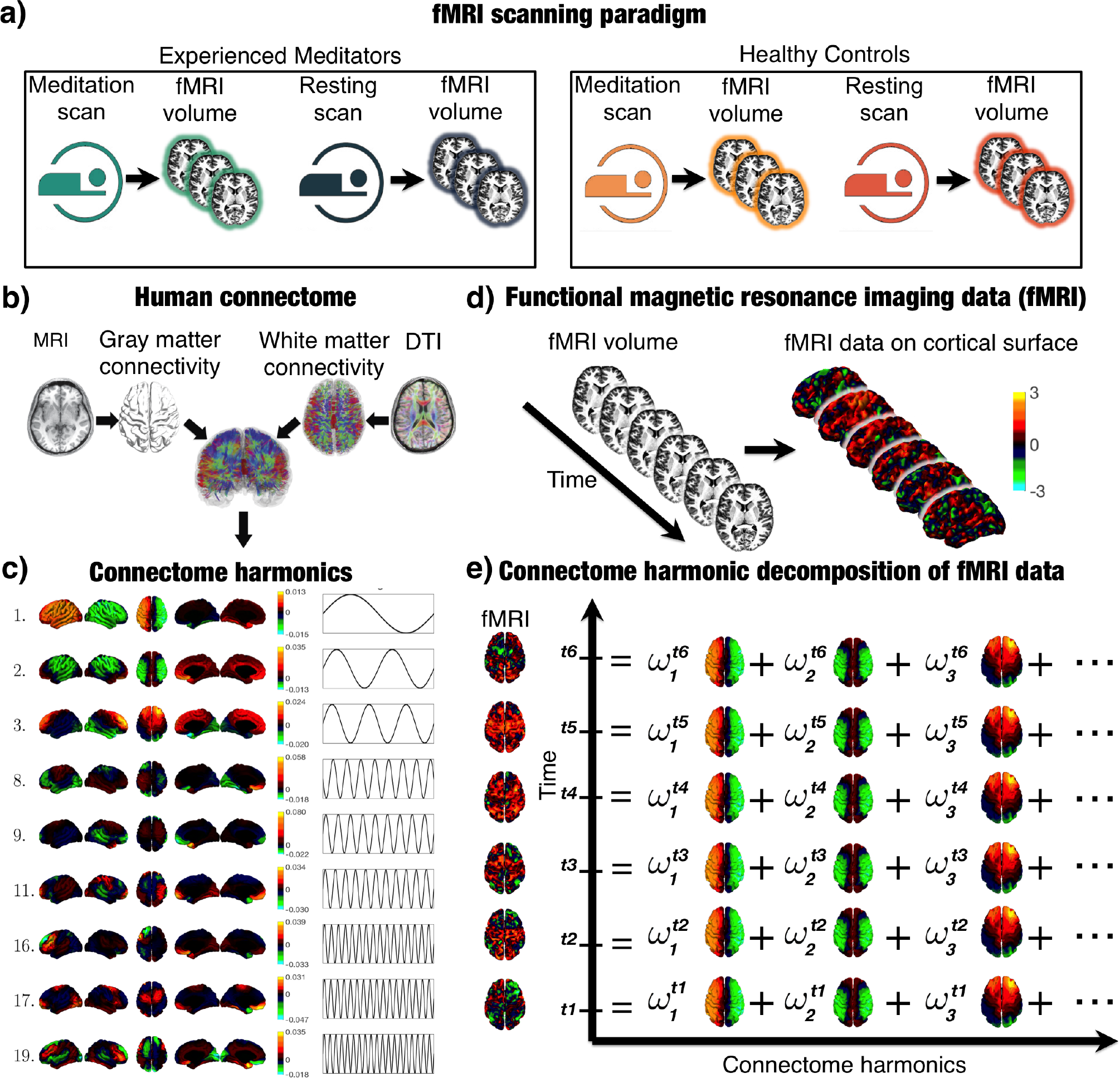
Illustration of the workflow. (**a**) demonstrates the fMRI scanning paradigm. T1 magnetic resonance imaging (MRI) data (**b**) is used to reconstruct the cortical surface between gray and white matter as shown in (**c**). Diffusion tensor imaging (DTI) data (**b**) is used to extract thalamo-cortical fibers using deterministic fiber tractography as shown in (**b**). Both, local and long-distance connections are combined to create the connectivity matrix of the human connectome as illustrated in (**b**). Connectome harmonics 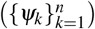 (**c**) are estimated by applying the Laplace operator ∆ to human connectome and computing its eigenvectors (∆*ψ*_*k*_ = *λ*_*k*_*ψ*_*k*_). Functional magnetic resonance imaging (fMRI) data (*ℱ*) as illustrated in (**d**) is decomposed in to the activation of connectome harmonics 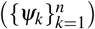 yielding the power of activation of each of these brain states for each time instance 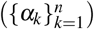 as delineated in (**e**).

### Mindfulness meditation increases the power, energy and complexity of brain activity

To study the immediate and long-term effects of mindfulness meditation, we applied the CHD to fMRI data acquired from two different groups (i.e. experienced meditators and healthy meditation-naive controls) in two different conditions (i.e. during meditation and during rest). For each fMRI condition, we decomposed the brain activity into connectome-specific waves and estimated the global effects in terms of power and energy across the whole connectome harmonic spectrum (see Methods). We found that the total power of brain activity was significantly higher in the meditator group during meditation compared to resting state (*p <* 10^−2^, two-sample t-test) (see Figure 2**a**). No significant difference was found in the control group between the two conditions (meditation vs. rest).

**Figure 2.**
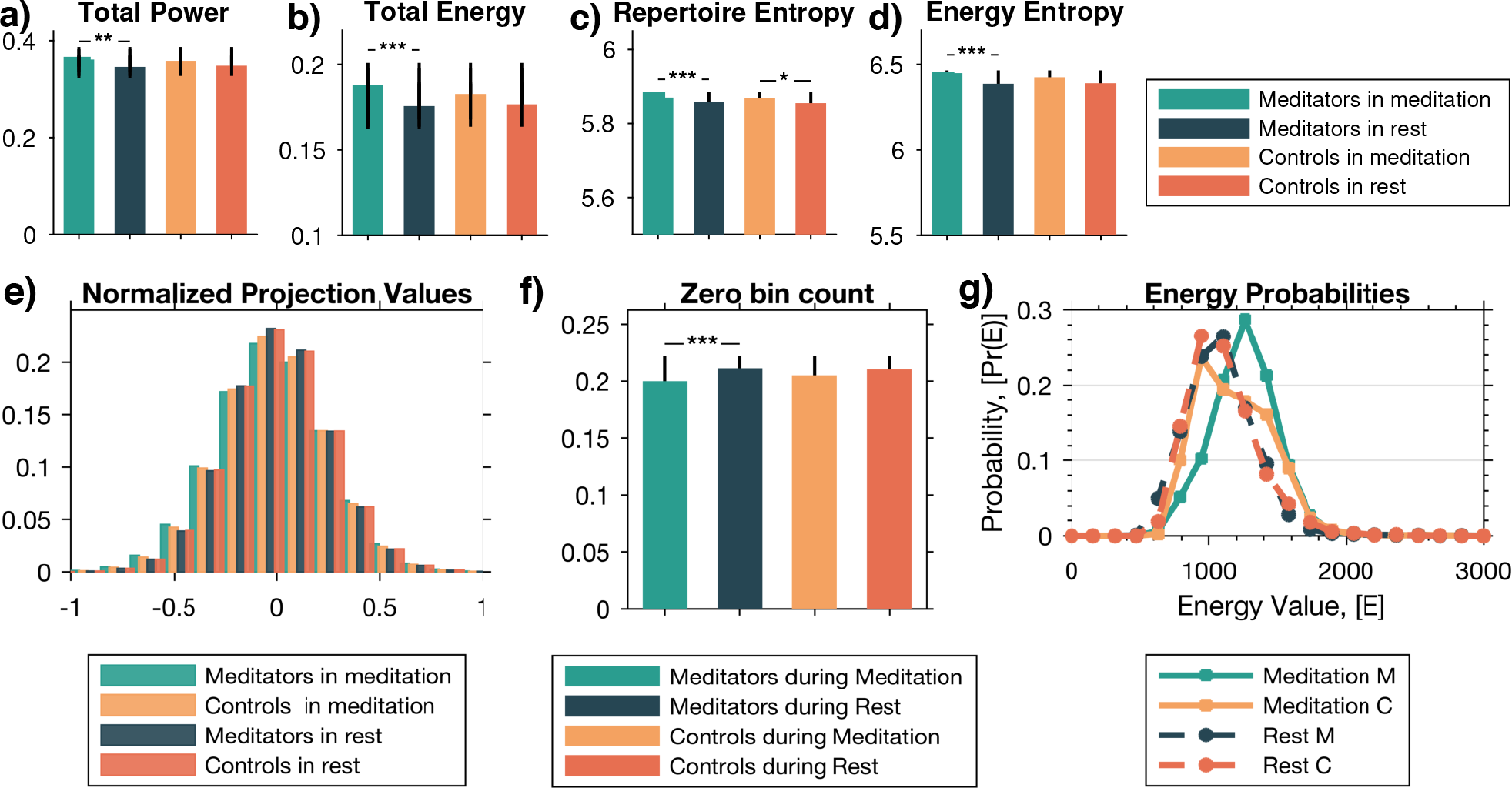
Total power (**a**), total energy (**b**), entropy of the complete power spectrum (**c**) and entropy of the complete energy spectrum (**d**) of all harmonic brain states for all 4 conditions, where stars indicate significant differences (∗ for p *<* 0.05, ∗∗ for *<* 10^2^ and ∗ ∗ ∗ for p *<* 10^3^, two-sample t-test) between each pair of meditation vs. rest conditions with indicated p-values. (**e**) Probability distribution of the occurrence of projection values (the amount of contribution) of connectome harmonics after normalization of each harmonic’s contribution by the maximum value of the baseline (rest for healthy control group) condition, shown for all 4 conditions. (**f**) The number of harmonics that have zero contribution to brain activity in each condition (meditation vs rest) in each group (meditators vs controls). (∗ ∗ ∗ for p *<* 10^3^, two-sample t-test). (**g**) Probability distribution of achieving different energy states for all 4 conditions.

Second, we estimated the total energy for each condition by combining the power of activation of each connectome harmonic by its intrinsic energy, averaged across all time points and subjects (see Methods). As the intrinsic energy of each connectome harmonic is closely related to its frequency, the energy can be seen as the frequency-weighted version of the power. We found that the total energy also significantly increases in the meditator group during meditation compared to rest (*p <* 10^−3^, two-sample t-test) (see Figure 2**b**), while again no significant difference was found for the control group.

To further evaluate the complexity of the power and energy spectra in each condition, we estimated the entropy of the complete power and energy spectrum of connectome harmonics in each condition yielding the measures of repertoire entropy and energy entropy, respectively. We found that the repertoire entropy (entropy of the power spectrum) significantly increased in both groups during meditation compared to resting state, yet the significance was stronger for the meditator group (*p <* 10^−3^, two-sample t-test) (see Figure 2**c**) compared to the meditation-naive control group (*p <* 0.05, two-sample t-test) (see Figure 2**c**).

This significance is diminished for the difference between resting state and meditation in the control group for the energy entropy measure, while it remained highly significant for the difference between resting and meditation conditions for experienced meditators (*p <* 10^−3^, two-sample t-test) (see Figure 2**d**).

These results suggest that mindfulness meditation significantly increases the energy, power and complexity of brain activity during meditation in experienced meditators.

### Mindfulness meditation expands the repertoire of harmonic brain states

To examine the repertoire of active connectome harmonics in each condition, we estimated the distribution of normalized projection values, where the normalization enables the comparison of distributions in different conditions. To this end, we compute the probability distribution of the occurrence of projection values (the amount of contribution) of connectome harmonics after normalization of each harmonic’s contribution by the maximum value of the baseline (rest for healthy control group) condition. As illustrated in (Figure 2**e**), the distribution widens during meditation for the experienced meditator group indicating an expansion of the repertoire of activated connectome harmonic brain states. To test the significance of this repertoire expansion, we evaluated the change in the bin corresponding to non-activated connectome harmonics (zero bin count) for all 4 conditions (Figure 2**f**). This evaluation revealed that the number of connectome harmonics which do not contribute to brain activity (i.e. non-activated) is significantly reduced for experienced meditators during meditation, whereas no significant difference was observed for the control group. There results demonstrate the significant expansion of the repertoire of active connectome harmonics during mindfulness meditation for experienced meditators.

Finally, to explore whether mindfulness meditation can lead to the obtainment of higher energy states, we evaluated the probability distribution of achieving different energy states in all 4 conditions. We found that the peak of the energy distribution shifted towards high energy states for experienced meditators during meditation (green curve in Figure 2**g**) compared to all other conditions. We observed a slight increase in the probability of high energy states for the control group during meditation (yellow curve in Figure 2**g**) compared to the resting state (orange curve in Figure 2**g**), yet this increase did not lead to a shift of the characteristic peak (the maximum) of the distribution towards high energy states, as it did for the experienced meditators.

These results reveal that mindfulness meditation leads to expanded repertoire of harmonic brain states for experienced meditators, which is also accompanied by a greater probability of achieving higher energy brain states.

### Mindfulness meditation alters harmonic brain states in a frequency-selective manner

In order to investigate weather meditation induced frequency-specific alterations on the harmonic brain states, we analysed the complete energy spectrum of connectome harmonics. Figure 3**a** illustrates the energy spectrum for the complete range of connectome harmonics binned into 15 bins in the logarithmic space as in previous studies^28,31^ for the experienced meditators and the control group during meditation and resting state.

**Figure 3.**
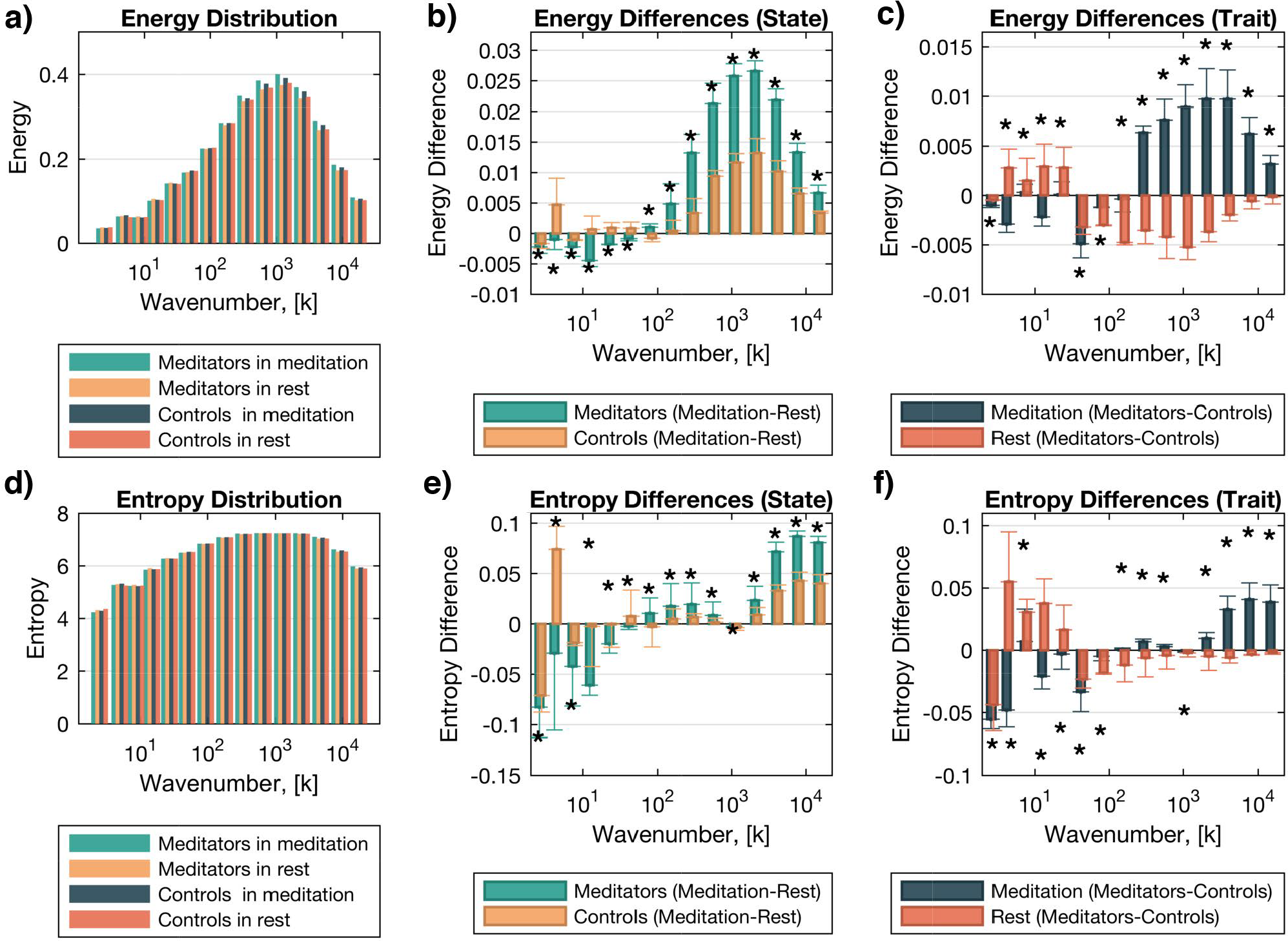
Energy spectrum (**a**) for the complete range of connectome harmonics binned into 15 bins in the logarithmic space as in previous studies^28, 31^ for the experienced meditators and the control group during meditation and resting state shown in green, dark blue, yellow and orange bars, respectively. (**b**) shows the energy differences between meditation and rest conditions for experienced meditators (green bars) and control group (yellow bars). (**c**) illustrates the energy differences between meditators and controls during meditation (dark blue bars) and in resting state (orange bars). Entropy of the energy spectrum (**d**) for the complete range of connectome harmonics binned into 15 bins in the logarithmic space. (**e**) shows the entropy differences between meditation and rest conditions for experienced meditators (green bars) and control group (yellow bars). (**f**) illustrates the entropy differences between meditators and controls during meditation (dark blue bars) and in resting state (orange bars).

To explore the frequency-specific alterations induced by meditation, we evaluated the differences in the connectome harmonics energy spectrum between meditation and rest conditions for both groups (Figure 3**b**). We found an increase in the connectome harmonic energy spectrum for a broad rage of high frequency harmonics (wave number larger than 200 out of 18715 connectome harmonics) in both groups, yet this increase was significantly larger in the experienced meditator group compared to healthy controls. Furthermore, we observed a significant decrease in the energy spectrum of connectome harmonics with wave number smaller than 200 (out of 18715 connectome harmonics) for the experienced meditator group, whereas we did not find the same effect for the control group. Interestingly, the energy differences of the complete connectome harmonic spectrum between meditation and rest for the experienced meditator group exhibited remarkably similar profile to that of the energy differences between the psychedelic state compared to placebo reported for various psychedelic compounds^28,31,33^. This similarity can be attributed to similar changes induced by the psychedelic state and meditation for experienced meditators caused by the less limited, more flexible exploration of brain state repertoire in these two states, as hypothesized by the entropic brain theory^34,35^.

In order to study the long-term effects of meditation in a frequency-specific manner, we analysed the groupwise differences between meditators and controls in the connectome harmonics energy spectrum for both conditions (Figure 3**c** dark blue bars). Once again we found an increase in the connectome harmonic energy spectrum for a broad rage of high frequency harmonics (wave number larger than 200 out of 18715 connectome harmonics) in meditation, in line with our previous analysis. This finding confirms that meditation leads to an increase in the energy of high frequency connectome harmonics and a suppression of the low frequency connectome harmonics and these changes in the energy profile of the connectome harmonic spectrum are significantly larger in the experienced meditator group compared to healthy controls. Remarkably, when evaluated for the resting state condition, the energy profile changes between the meditator and control group reversed (Figure 3**c** orange bars) compared to that of the meditation condition (Figure 3**c** dark blue bars). This result demonstrates the long-term effect of meditation on the resting state brain and indicates that experienced meditators exhibit the inverse effects in terms of frequency-specific energy changes in their brain dynamics during resting state compared to meditation. This crucial finding can be attributed to calmer brain dynamics that experienced meditators may experience in resting state as a result of their regular meditation practice.

In order to test this hypothesis, we evaluated the entropy of the energy spectrum for all conditions as illustrated in Figure 3**d**. Once again, we studied the differences between conditions (meditation and rest, (Figure 3**e**) and between groups (meditators and controls, (Figure 3**f**), this time in terms of entropy of the connectome harmonic energy spectrum. We found that the entropy changes in condition-wise (Figure 3**e**) and group-wise (Figure 3**f**) evaluations showed similar characteristics to the energy changes in condition-wise (Figure 3**b**) and group-wise (Figure 3**c**), respectively, even though the exact entropy profiles were slightly different than those of energy profiles. These results confirm that the energy changes in connectome harmonic spectrum are accompanied by the changes in complexity of brain activity. Specifically, we observed that the complexity of brain dynamics attributed to low frequency connectome harmonics showed significant reduction during meditation for experienced meditators while the complexity of brain dynamics corresponding to a broad range of high frequency connectome harmonics significantly increased (Figure 3**e**). These changes were observed for the experienced meditator group only indicating that the differences in the complexity of brain dynamics only occurs during meditation for experienced meditators and not for the first time meditators. Finally, we found that the entropy changes between groups (meditators and controls) in meditation and resting conditions was in line with the energy changes confirming that the long term effects of meditation leading to changes in connectome harmonic energy spectrum are accompanied by alterations in complexity of brain dynamics in the corresponding energy bands (Figure 3**f**). Crucially, this finding suggests that the brain activity of experienced meditators exhibits higher complexity during meditation and a lower complexity during resting state supporting the hypothesis that the meditators experience calmer brain dynamics during resting state.

### State and trait changes in the connectome harmonics energy and complexity spectra for long-term mindfulness meditators

There has been a long discussion in the meditation science literature about changes in brain states during a meditation session versus ‘trait’ changes seen in the brains of long-term meditators during a resting (non-meditation) condition^36–40^. The above analysis, specifically the energy and complexity spectra shown in Figures 3**c** and 3**f**, respectively, essentially reveals the signature of the meditation state and the resting trait of long-term meditators. These results reveal that during the meditation ‘state’ the brain activity of long-term meditators, relative to controls, exhibits an energy and entropy/complexity decrease in the low frequency harmonics up to a wavenumber of 200 and an increase of energy and complexity of the higher frequency harmonics. In contrast, during the resting condition the brain activity of long-term meditators, relative to controls, shows an energy and entropy/complexity increase in the lower frequency harmonics up to a wavenumber of about 30 (but excluding the zero-th (constant) harmonic which we will discuss below) and an decrease in the higher frequency harmonics, uncovering the long-term meditator ‘trait’ signature.

Furthermore, the energy and entropy of the first connectome harmonic, representing the basic oscillation between the left and right cerebral hemispheres, decreases in both the meditation state and the resting ‘trait’, relative to controls. This contrast of state and trait signatures is reminiscent of the training effect of aerobic exercise where during the exercise ‘state’ the heart rate increases, yet the long-term exercisers are shown to have lower resting ‘trait’ heart rates. Analogously, long-term meditators display higher complexity and energy in the high frequency harmonics during meditation but lower than controls complexity and energy in the high frequency harmonics in a resting state (the ‘trait’). The converse is found for the low frequency harmonics: lower complexity and energy during meditation but higher than controls complexity and energy in a resting state (the ‘trait’). Remarkably, these results are also in agreement with the recent findings of the review^41^, which reveals that neural activity during the meditative state has a higher complexity when compared to waking rest or mind-wandering, and the brain activity of experienced meditators as a trait exhibits decreased baseline complexity when compared to novices and controls.

More parallels with the previous literature emerge when we focus on the studies exploring inter- and intra-network connectivity changes due to meditation. In previous work, we have suggested that the low frequency harmonics are associated with the large-scale resting-state cortical networks^30^. There is now increasing convergent evidence that trait mindfulness is associated with changes in functional connectivity within and between resting-state networks related to attention and cognition such as the default mode network (DMN), the salience network (SN) and the central executive network (CEN)^18,26,42–44^. It is thus suggestive that the connectome harmonic signatures of meditation and rest in terms of energy and complexity spectra of long-term meditators indicate a training effect, where during meditation a disruption of patterns of functional connectivity between the DMN, CEN and SN occurs, which then leads to new patterns of enhanced functional connectivity between these networks during rest. Hence, our findings suggest the first explanatory mechanism for the occurrence of these inter- and intra-network connectivity observed state and trait changes in the meditative brain.

### Mindfulness meditation decreases cross-frequency correlations across connectome harmonics

In order to understand the mechanisms underlying the complexity changes in connectome harmonic spectrum during meditation for experienced meditators, we explored the coupling (correlation) of the activation profiles of different frequency harmonics over time. To this end, we evaluated the percentage of occurrence of different cross-frequency correlation values after taking the histogram of the cross-frequency correlation matrix using 20, 100 and 1000 bins as shown in Figure 4**a**, 4**b** and 4**c** respectively. Independent of the histogram bin number that is used, we found that the cross-correlation values showed a significant reduction (indicated in the higher zero peak) for experienced meditators during meditation compared to all other three conditions. This finding not only confirms that brain activity of experienced meditators during meditation exhibits higher complexity but also reveals that this increase in complexity is accompanied by the reduced cross-frequency coupling (hence higher degrees of freedom) across different frequency connectome harmonics.

**Figure 4.**
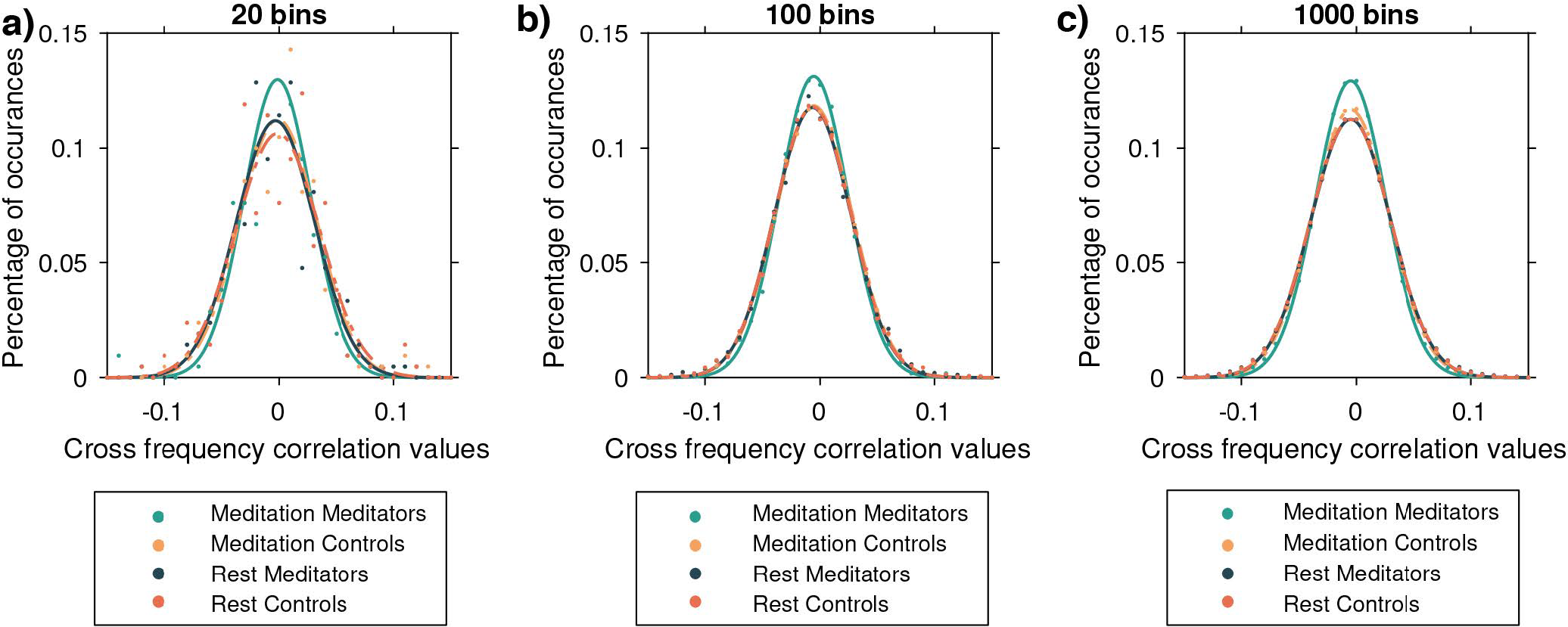
The percentage of occurrence of different cross-frequency correlation values after taking the histogram of the cross-frequency correlation matrix using (**a**) 20, (**b**) 100 and (**c**) 1000 bins.

### Correlation between subjective measures and energy changes

Finally, we tested weather the energy and entropy changes in connectome harmonic spectrum correlated with the subjective measures of hours of meditation experience, age and years of education. To this end, we evaluated the correlations of the subjective measures with the average energy changes in the whole connectome harmonic spectrum (Figure 5**a**-**c**) and with the entropy of the connectome harmonic energy spectrum (Figure 5**d**-**e**). We found that the entropy of the energy profile showed near significant correlation with the hours of meditation experience (Figure 5**d**, p *<* 0.057) suggesting that the complexity of brain dynamics during meditation correlates with the hours of experience; i.e. more experienced meditators exhibit more complex brain dynamics during meditation. This significance was not found in the correlations between hours of meditation experience and the average energy changes (Figure 5**a**, p *<* 0.081). No significant correlation was found between the age of the experienced meditators and average energy changes (Figure 5**b**) or entropy changes (Figure 5**e**) of the connectome harmonic energy spectrum. Intriguingly, the years of education of the meditators showed significant correlations with both, average energy changes (Figure 5**c**, p *<* 0.021) and the the entropy changes (Figure 5**f**, p *<* 0.017) of the connectome harmonic energy spectrum. These findings suggest that years of education alters brain dynamics in a manner that is reflected in both the average energy changes as well as the complexity of the connectome harmonic energy spectrum, whereas meditation induces more specific alterations that only link to the complexity of the connectome harmonic activation patterns and is not predictable by a more simple measure such as the average energy changes in the connectome harmonic spectrum.

**Figure 5.**
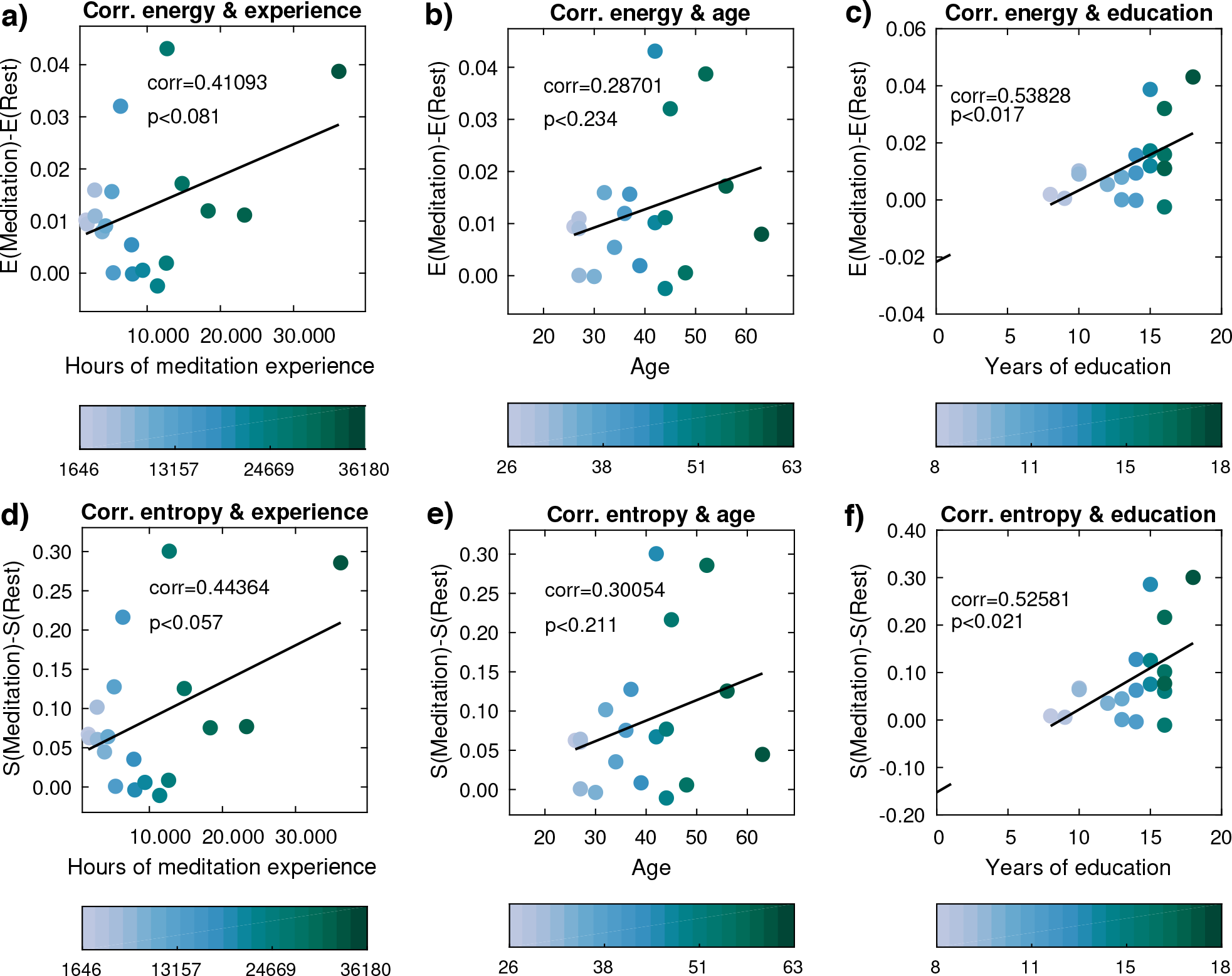
Correlations of the subjective measures of (**a**) hours of experience, (**b**) age and (**c**) years of education with the average energy changes in the whole connectome harmonic spectrum, respectively. Correlations of the subjective measures of (**d**) hours of experience, (**e**) age and (**f**) years of education with the entropy of the connectome harmonic energy spectrum, respectively.

## Discussion

In this paper we reveal both the immediate (state) and long-term (trait) effects of meditation on whole-brain dynamics by decomposing fMRI data of meditation-na*ï*ve healthy participants and experienced meditators in to their harmonic signatures. To this end, we use the connectome harmonic decomposition framework^28–30^, which represents brain activity in terms of the frequency-specific harmonic brain states defined by connectome harmonics^30^. Previous research demonstrated that these harmonic waves emerge from the interplay between excitatory and inhibitory brain activity in the human brain and these harmonic waves termed “connectome harmonics” significantly match the well-known functional networks of the human brain^30^. Importantly, by definition connectome harmonics extend the well-known Fourier basis to the specific structure of the brain, introducing a new harmonic language in which any pattern of brain activity can be expressed as a weighted combination of connectome harmonics. This allows for a spatial harmonic decomposition of brain activity to study cortical dynamics^28,29^. Furthermore, the change of representation of brain activity to the activation of frequency-specific brain states (i.e. connectome harmonics) provides a paradigm shift in the understanding of how different frequency brain states relate to the phenomenological experience.

Versions of this harmonic decomposition framework have been successfully utilised to decode the harmonic signatures of various mental states including psychedelic-induced altered states of consciousness^28,33^, propofol-induced loss of consciousness^31^, vegetative state and minimally conscious state^31^. In this work, we apply this framework for the first time to fMRI data of experienced meditators and healthy, meditation-naive controls acquired during meditation and resting state.

Our results reveal that mindfulness meditation increases the power, energy and complexity of brain activity during meditation for experienced meditators but not for healthy controls. Furthermore experienced meditators are found to achieve higher energy states during meditation compared to resting state and compared to both conditions (meditation and rest) of the control groups. The observed increase in the complexity of brain dynamics is found to be linked to the expansion of the activated connectome harmonics repertoire as well as to the decreased cross-frequency coupling across different connectome harmonics.

The meditation-induced changes in the connectome harmonic energy spectrum are found to be frequency specific, where experienced meditators exhibit a decrease in the energy of low frequency connectome harmonics accompanied by an increase in the energy of a wide range of high frequency connectome harmonics. Remarkably, previous research has reported a similar energy profile in the psychedelic-induced altered state of consciousness^28,33^ and the exact opposite energy distribution pattern in the propofol-induced loss of consciousness^31^ and in vegetative state^31^. Taken together these findings suggest that the combined effect of low frequency suppression and high frequency amplification can be the characteristic signature of an expanded state of consciousness experienced during meditation (for experienced meditators) and psychedelic-induced altered state, whereas the opposite profile is characteristic of a rather limited state of consciousness experienced during anestesia-induced loss of consciousness and vegetative state. It is worth to note that the group-wise difference between experienced meditators and meditation-naive healthy controls exhibits inverse energy profiles in their brain dynamics during resting state compared to meditation. This finding indicates that meditators experience calmer brain dynamics in resting state as a result of their regular meditation practice, which is also confirmed by our complexity analysis.

In particular, the frequency-specific changes in brain dynamics of experienced meditators in terms of both measures, energy and entropy, display the inverse profile in resting state, whereas the same effect is not found for meditation-naive control groups. These findings reveal for the first time the immediate (state) as well as the long-term (trait) signatures of the meditative brain in terms of energy and complexity of the connectome harmonic spectrum.

Besides the frequency-specific alterations in the brain activity of experienced meditators during meditation, our results revealed an expanded repertoire of harmonic brain states is attained by the brain activity of experienced meditators during meditation. This repertoire expansion is found to be also accompanied by an increased complexity of brain dynamics as reflected in the reduced cross-frequency coupling (hence higher degrees of freedom) across different frequency connectome harmonics. Finally, the specific changes in the increased complexity and energy of the complete connectome harmonic spectrum showed nearly significant correlations between the hours of meditation experience and significant correlations with the years of education of experienced meditators. These correlations emphasize the functional role of the energy of complexity changes of the connectome harmonic spectrum and demonstrate the usefulness of the connectome harmonic decomposition to link brain activity to subjective measures and experience.

## Methods

### Participants

A total of 20 experienced meditators and 20 healthy controls participated in this study. All of them had no history of past neurological disorders. The meditator group was recruited from the Vipassana communities of Barcelona, Catalonia. Meditators had more than 1,000 hours of meditation experience and maintained daily practice (*>* 1 hour/day) (7 females; mean age=39.8 years; SD=10.29 years; education=13,6 years; and meditation experience=9.526,9 hours; SD=8.619,8 hours). Healthy controls had no previous experience in meditation practice and were well-matched for age, gender, and educational level (7 females; mean age= 39,75 years; SD=10,13 years; education=13,8 years). No significant differences in terms of age, educational level and gender were found between groups. One meditator was removed before the preprocessing of the data due to incidental imaging findings in the MRI session and, as explained in the Preprocessing section, 4 participants were excluded due to movement artifacts during the MRI session. Therefore, the final sample of the study included: 19 meditators during wakefulness and meditation, 16 controls during wakefulness and during meditation. The dataset has been already published and explained in^45^. The experimental protocol for this study was approved by the Bellvitge Hospital Ethics Committee by following the Helsinki Declaration. Participants provided written consent before started the study and were compensated for their participation.

### Experimental Procedure

Approximately a total of 15 minutes in both conditions (resting-state and meditation), were analyzed. During resting-state, participants were asked to stay as motionless as possible, relaxing and not thinking about anything in particular while looking at a cross fixated on the screen. After the resting-state acquisition, participants underwent the focused attention meditation task. Meditators were asked to practice the mindfulness of breathing (i.e., anapanasati in Pali). In this form of meditation, subjects have to be aware of the sensations in the area of the nostrils produced by natural breathing. If a thought arises or they fall into a distraction, they have to recognize it and come back to natural breathing without judgment. Controls were instructed in anapanasati meditation before the fMRI scan. They confirmed that they had understood the task after having done a simulation.

### MRI Data Acquisition

MRI images were acquired through a 3T whole-body Siemens TRIO scanner (Hospital Clínic, Barcelona) using a 32-channel receiver coil. The high resolution T1-weighted images were acquired with repetition time (TR) of 1970ms; echo time (TE) of 2.34ms; inversion time (IT) of 1050ms; 208 sagittal slices; flip angle=9 degrees; field of view (FOV) = 256mm; and isotropic voxel size 1x1x1mm with no gap between slices. Functional images were acquired by a single shot gradient-echo EPI sequence (TR = 2000ms; TE = 29ms, FOV = 240mm, in-plane resolution 3mm, 32 transversal slices thickness = 4mm, no gap between slices, and flip angle = 80 degrees).

### fMRI Pre-processing

The resting-state and meditation fMRI data were pre-processed using the Data Processing Assistant for Resting-State fMRI (DPARSF)^46^ based on Statistical Parametric Mapping (SPM12) http://www.fil.ion.ucl.ac.uk/spm. A total of 450 volumes in each condition were analyzed. T1 and EPI images were manually re-oriented. Pre-processing steps included: 1) discarding the first 10 volumes to avoid MRI saturation effects; 2) slice-timing correction; 3) realignment for motion correction; T1 co-registration to functional image; 5) European regularisation segmentation; 6) removal of spurious variance through linear regression: six parameters from the head-motion correction, the global mean signal, the white matter signal (WM), and the cerebrospinal signal (CSF), CompCor^47^; 7) linear trend removal; 8) spatial normalization to the Montreal Neurological Institute (MNI); 9) Gaussian Kernel smoothing of 6-mm FWHM; and 10) band-pass filtering (0.01-0.25 Hz)^48,49^. Moreover, we calculated the frame-wise displacement (FD) of head motion as previously described by Jenkinson and colleagues^50^. Participants with mean FD grater than 2 standard deviations above the mean motion group were excluded from the analysis^51^. This resulted in the exclusion of 4 controls during meditation and 1 control during rest due to head motion.

### Computation of connectome harmonics

Connectome harmonics were calculated by using a sample of 10 unrelated subjects (6 female, age between 22 and 35 years), as successfully performed in previous studies^52,53^. DTI and T1-weighted structural images were obtained and made available by the WU-Minn Human Connectome Project (1U54MH091657), funded by the 16 NIH Institutes and Centers that support the NIH Blueprint for Neuroscience Research; and by the McDonnell Center for Systems Neuroscience at Washington University. Datasets were preprocessed following the minimal preprocessing pipelines for structural and diffusion MRI of the Human Connectome Project^54^. In brief, Freesurfer http://freesurfer.net was used to reconstruct the cortical surfaces. The T1-weighted image was segmented to obtain the white and grey matter independently for each subject. Then, deterministic tractography was applied to estimate the corticocortical and thalamocortical white matter fibres using the MATLAB implementation of Vista Lab, Stanford University http://white.stanford.edu/newlm/index.php/MrDiffusion. The DTI and the cortical surface were registered for each subject. Centred around each vertex (node) -in total 20,484-eight seeds were initialized and the deterministic tractography was estimated using the following parameters: fractional anisotropy (FA) threshold 0.3 (i.e., FA*<*0.3 being termination criteria for the tracking), minimum tract length 20 mm, and the maximum angle between two contiguous tracking steps 30°.

The graph of the human connectome is described as *𝒢* = (*𝒱, ℰ*), where 𝒱 are the vertices sampled from the gray matter surface by the nodes *𝒱* = {*ν*_*i*_, | *i* ∈ 1, …, *n*} (with *n* = 20.484), and *ℰ* are the edges representing the estimated tractography connections between the vertices *ℰ* = {*e*_*i j*_, | (*ν*_*i*_, *ν*_*j*_) ∈ *𝒱 x 𝒱* }. For each participant, an adjacency matrix was computed as:

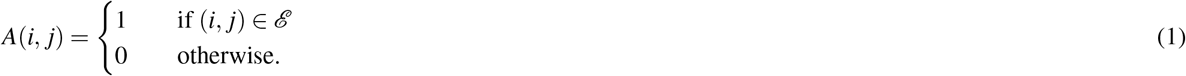

The resulting adjacency matrices were then averaged yielding an average structural connectivity across subjects *Ā* (i.e., the group average adjacency matrix). Then, the symmetric graph Laplacian ∆_*𝒢*_ was calculated on the average connectome graph to estimate the discrete counterpart of the Laplace operator ∆ applied to the human connectome (i.e., the connectome Laplacian) as:

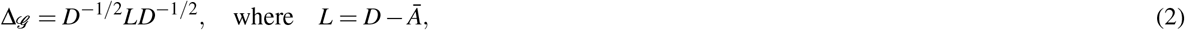

and D represents the matrix degree of the graph as:

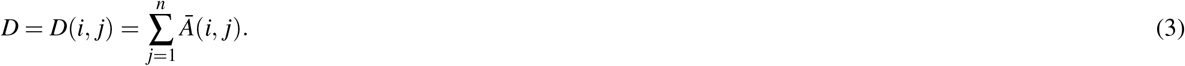

Finally, the connectome harmonics *ψ*_*k*_, *k* ∈ {1,…, *n*} were calculated by applying the following equation:

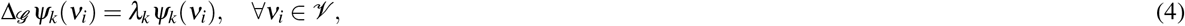

where *λ*_*k*_, *k* ∈ {1,…, *n*} represents the eigenvalues of ∆ _*𝒢*_ .

### Connectome Harmonic Decomposition of Meditation and Resting-State fMRI

The meditation and resting-state fMRI data were then rendered onto the cortical connectome harmonics coordinates by applying the -volume-to-surface-mapping command of the Human Connectome Project Workbench. This registration yields the time course *ℱ* (*v, t*) for all vertices *ν* ∈ *𝒱* on the cortex. Then, the spatial cortical activity pattern *ℱ t*_*i*_(*v*) for each time point *t*_*i*_ ∈ {1, …, *T* } (with *T* = 440) of the time course *ℱ* (*v, t*) was decomposed into the activity of connectome harmonics 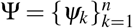 as:

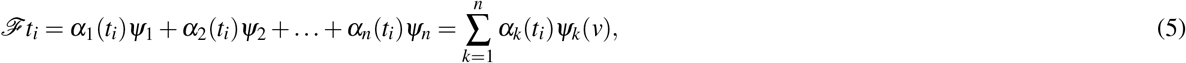

where the temporal activity *α*_*k*_(*t*) of each connectome harmonic *ψ*_*k*_ was calculated by rendering the fMRI data *ℱ* (*v, t*) onto that particular harmonic. The coefficients *α*_*k*_ were given by:

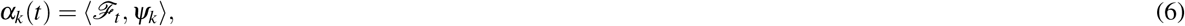

where the subscripts *k* and *t* corresponds to the connectome harmonic wavenumber *ψ*_*k*_ and the time point, respectively.

### Power and energy of brain states

The activation power of each connectome harmonic *ψ*_*k*_, *k*∈ *{*1, …, *n}* at each time point *t* for each fMRI condition was computed as the activation strength for each connectome harmonic *ψ*_*k*_ as:

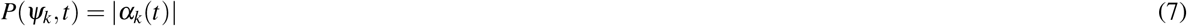

and the total power of a harmonic brain state at a given time point *t* was given by:

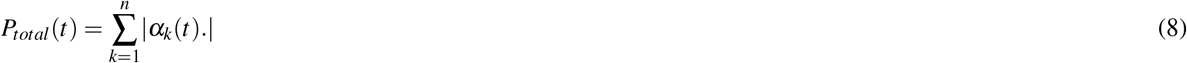

The energy of each connectome harmonic *ψ*_*k*_, *k* ∈*{*1, …, *n}* in the cortical activity pattern at a particular time point *t* for each fMRI condition was computed by combining the total activation strength of each connectome harmonic with its intrinsic energy which is given by 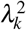 (Equation 4). Thus, the energy of a brain state was defined *ψ*_*k*_ as:

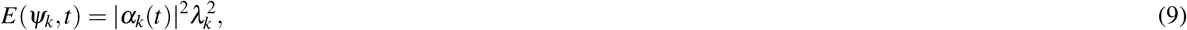

and the total energy of a harmonic brain state at a given time point *t* was given by:

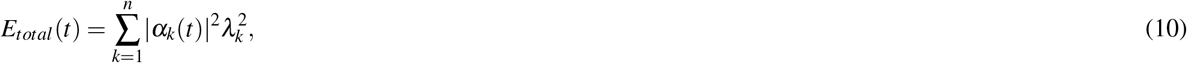

where the equation can be rewritten as:

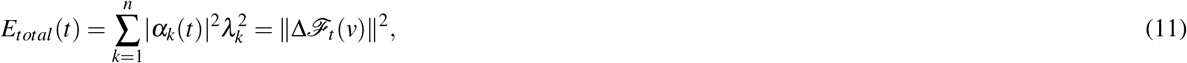

where the Laplace operator ∆ obtains the total flow activity. That is, the total energy of brain activity corresponds to the total flow of neural activity across the cortex at a given time *t*. The total power and energy of each brain state are given by the sum across all time points. The power corresponds to the product between a connectome harmonic and the cortical activity pattern at a given time. Given that the connectome harmonics are orthonormal (i.e., ‖*ψ*_*k*_‖= 1, ∀ *k*), the lower bound of the power is 0, while the upper bound is determined by the cortical activity pattern. For energy values, the square of power is weighted by the square of each connectome harmonic eigenvalue, and the eigenvalues are delimited by *λ*_*k*_ *<* [0, …, 2] for the connectome Laplacian.

### Cross-frequency correlations between brain states

Cross-frequency correlations were obtained by computing the Pearson’s linear correlation coefficient *r* between each pair of brain states (*ψ*_*i*_, *ψ*_*j*_) as:

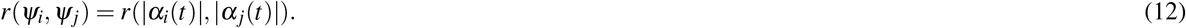

## Acknowledgements (not compulsory)

Acknowledgements should be brief, and should not include thanks to anonymous referees and editors, or effusive comments. Grant or contribution numbers may be acknowledged.

## Author contributions statement

Must include all authors, identified by initials, for example: A.A. conceived the experiment(s), A.A. and B.A. conducted the experiment(s), C.A. and D.A. analysed the results. All authors reviewed the manuscript.

## Additional information

To include, in this order: **Accession codes** (where applicable); **Competing financial interests** (mandatory statement).

The corresponding author is responsible for submitting a competing financial interests statement on behalf of all authors of the paper. This statement must be included in the submitted article file.

